# snPATHO-seq: unlocking the FFPE archives for single nucleus RNA profiling

**DOI:** 10.1101/2022.08.23.505054

**Authors:** Andres F Vallejo, Kate Harvey, Taopeng Wang, Kellie Wise, Lisa M. Butler, Jose Polo, Jasmine Plummer, Alex Swarbrick, Luciano G Martelotto

## Abstract

FFPE (formalin-fixed, paraffin-embedded) tissue archives are the largest repository of clinically annotated human specimens. Despite numerous advances in technology, current methods for sequencing of FFPE-fixed single-cells are slow, labour intensive, insufficiently sensitive and have a low resolution, making it difficult to fully exploit their enormous research and clinical potential. Here we introduce single nuclei pathology sequencing (snPATHO-Seq), a sensitive and efficient high-throughput platform to profile the transcriptome of single nuclei extracted from formalin-fixed paraffin-embedded (FFPE) samples. snPATHO-Seq combines an optimised nuclei extraction protocol from archival samples with 10x Genomics probe-based technology targeting the whole transcriptome. We performed direct comparison of the Fixed RNA Profiling (FRP) and established 3’ single cell RNA-Sequencing (scRNA-Seq) workflows through a comprehensive bioinformatics analysis of matched fresh and fixed samples derived from the LNCaP prostate cancer cell line. FRP detected 2.1 times more transcripts in the fixed sample than the 3’ kit did in the fresh sample. Low mitochondrial genes detection using the FRP was translated into 99.9 percent of cells passing the QC filters, compared to 81.6 percent of cells using the v3.1 chemistry. We then optimized snPATHO-Seq and applied it to a human breast cancer metastasis to the liver collected at autopsy and preserved in FFPE, a particularly challenging sample type. Remarkably, at 28,000 reads/cell snPATHO-Seq was able to detect a median of 1850 genes/cell and 3,216 UMI counts/cell. Comparison of snPATHO-Seq with spatial transcriptomics data (10x Genomics Visium FFPE v1) derived from an adjacent section of the same sample revealed a strong correlation, validating the accuracy of the snPATHO-Seq data. Gene expression data from snPATHO-Seq was used to predict cell type composition within each spatial transcriptomic location via deconvolution. Overall, snPATHO-Seq enables high quality and sensitivity snRNA-Seq from preserved tissue samples, unlocking the vast archives of FFPE tissues and thereby allowing extensive retrospective clinical genomic studies.

## Introduction

Simultaneous sample processing of recently isolated cells is the most desirable approach to increase the data quality and prevent technical batch effects in scRNA-Seq experiments. However, fresh/frozen specimen procurement is not a standard clinical and diagnostic practise in most institutions, and fresh/frozen samples cannot be obtained for certain sample types. For diagnostic purposes, the vast majority of human tumour tissue is routinely formalin-fixed paraffin-embedded (FFPE). The estimated millions of FFPE tissue samples available provide a vast resource for the identification of disease pathways, biomarkers, and drug targets (Hester et al., 2016; Klopfleisch et al., 2011).

Molecular analysis of formalin-fixed paraffin-embedded (FFPE) specimens is difficult, because formalin fixation introduces several types of artefacts, primarily caused by protein and nucleic acid cross-link (Gilbert et al., 2007). Despite the fact that whole-exome and targeted sequencing analyses have been performed successfully on DNA extracted from FFPE tumour bulk samples (Basile et al., 2021; Fischer et al., 2022; Jang et al., 2021; Mehine et al., 2020) scRNA-Seq methods for transcriptomic investigations of FFPE tissue samples are still undeveloped. Previously, we developed a method for whole-genome single-cell copy number analysis of formalin-fixed paraffin-embedded samples. This method resulted in a single-cell copy map of breast cancer tissue that revealed the transition from in situ to invasive breast cancer, demonstrating significant potential for clinical cancer research (Martelotto et al., 2017). In this pre-print we provide the first glimpse of snPATHO-Seq, a sensitive and efficient high-throughput method to extract and sequence nuclei from FFPE, paving the way for clinical applications. We report a side-by-side comparison between the Fixed RNA Profiling kit and standard Gene Expression Profiling scRNA-seq (10x Genomics) to validate the robustness of the former as an alternative sequencing strategy in most cancer research laboratories. We then optimized and applied snPATHO-Seq to breast cancer samples; here we present data on a 4 yo FFPE a breast cancer metastasis to the liver collected at autopsy. Gene expression data from snPATHO-Seq was used for mapping the detected cell types into spatial transcriptomic data for producing spatial maps of cell types via deconvolution. We provided evidence to show that snPATHO-Seq is a reliable platform for analysing the transcriptomic profiles of FFPE tumor tissues, which has the potential to unlock the largely untapped cancer archives of biological material found in pathology archives. This, thereby, will likely enable extensive retrospective clinical genomic studies. snPATHO-Seq full protocol (**snPATHO-Seq Protocol**) is available in this manuscript supplementary material, hoping you can contribute information about to the robustness of the protocol.

## Material and Methods

### Patient material, ethics and consent for publication

The breast cancer metastasis to the liver collected at autopsy sample used in this study was collected with written informed consent under the SVH 17/173 protocol with approval from St Vincent’s Hospital Ethics Committee. Consent included the use of all de-identified patient data for publication. Samples were collected during an autopsy that commenced 4hrs after death. Tumour tissue was fixed in 10% neutral buffered formalin for 24 h and then processed for paraffin embedding. Sections were stained with hematoxylin and eosin (H&E) for standard histological analysis. The prostate cancer cell line LNCaP (American Type Culture Collection) was used for validating the fixation kit.

### LNCaP cell line processing

Cells growing in RPMI 1640 media containing penicillin (100 units/ml), streptomycin (100 μg/ml), and 10% FBS at 37°C in a 5% CO2 incubator were trypsinized and washed 3 times with PBS. Cells were split in two aliquots, one aliquot was used for standard, poly-A based, Gene expression profiling (see below) and the other was fixed using the Fixation of Cells & Nuclei for Chromium Fixed RNA Profiling kit (see below) following the user guide recommendations.

### Cell line standard single-cell RNA profiling and Sequencing

Standard single-cell gene expression libraries (polyA-based) were performed using the Chromium NextGEM Single-cell 3’ Reagent kit (v3.1, 10x Genomics) following user guide recommendations (CG000204 - Rev D) to capture and profile ∼5000 cells. Libraries were sequenced on a NextSeq500 (Illumina) following 10x Genomics’ recommendations.

### Cell line Fixation, FRP profiling and Sequencing

Cells were fixed and profiled following the user guide recommendations for Fixation of Cells & Nuclei for Chromium Fixed RNA Profiling (CG000478 - Rev A, 10x Genomics) to capture and profile ∼5000 cells/sample. Cells were counted using LUNA-FX7 cell counter (AO/PI viability kit, Logos). Libraries were constructed using the Chromium Fixed RNA Profiling CG000477 - RevB) as singleplex using the BC1 probe-set, incubating for 20 h at 42°C and using 12 cycles for Indexing PCR. Libraries were sequenced on a NextSeq500 (Illumina) following 10x Genomics’ recommendations.

### Nuclei suspension preparation for snPATHO-Seq

1-2 >25 μm-thick sections were washed three times with 1 mL Xylene for 10’ to remove the paraffin. Sample rehydration was done in sequential immersions in 1 mL ethanol for 1’ (2× 100%, followed by 1× 70%, 50% and 30% ethanol). The sample was inspected visually to ensure the complete removal of paraffin. The sections were then washed 3 times (2× 1 mL wash and 1× 800 μL final wash) with 1× PBS + 0.5 mM CaCl2 removing as much liquid as possible. The tissue was digested for 45-60’ at 37°C in a Thermomixer at 800 rpm in 1 mL of 1× PBS + 0.5 mM CaCl_2_ + 250 μg/mL Liberase (Roche) + 2 mg/mL of Collagenase (Sigma-Aldrich) + 1 U/mL RNAse Inhibitor. After the incubation, 400 μL of Ez Lysis Buffer (Sigma-Aldrich) was added to the sample, mixing by inversion 5× and centrifuged for 5’ at 850xRCF at 4°C. The pellets (released nucs and undigested tissue) were resuspended in 250 μL EzLysis buffer + 2%BSA + 1 U/mL RNAse Inhibitor and homogenized using a douncer/pestle by stroking 20 times. After homogenization, 750 uL EzLysis buffer + 2% BSA + 1 U/mL RNAse Inhibitor was added to the sample and continue disaggregating by pipetting using a P1000 pipette (10 times). Incubate on ice for 10’. After 10’ sample was passed through a 25 G needle for 20-30 times and filtered through 70 μm filter (pluriSelect). The sample was centrifuged for 5’ at 850xRCF at 4°C and washed with 800 μL of EzLysis buffer and pelleted once more before resuspending the nuclei in 500 μL of 1x Fix & Perm Buffer (PN-2000517) for 1 h at RT. After this step, the sample were passed through a 40 μm PluriStrainer filter (not Flowmi!) and pelleted for 5’ at 850xRCF at 4°C. The nuclei were then pelleted 5’ at 850xRCF at 4°C, washed twice with PBS 0.5x + 0.02% BSA and resuspend in 500-1000 μL of PBS 0.5x + 0.02% BSA and rested on ice. Nuclei was counted using LUNA-FX7 cell counter (AO/PI viability kit, Logos). A detailed protocol for nuclei preparation for snPATHO-Seq is provided as supplementary material. Nuclei library was constructed using the Chromium Fixed RNA Profiling CG000477 - RevB) as singleplex using the BC1 probe-set, incubating for 20 h at 42°C and using 14 cycles for Indexing PCR.

### Spatial transcriptomics

A 5 μm thick section was prepared from the FFPE tissue block and processed using the Visium Spatial Gene Expression for FFPE Kit v1 (10x Genomics) according to the manufacturer’s instructions. Briefly, sections were stained with H&E and imaged followed by probe hybridisation and ligation. The captured probe library was then quality controlled and sequenced using an Illumina NovaSeq 6000 system for 28, 10, 10 and 50 cycles for Read 1, i7, i5 and Read 2 sequences respectively.

Reads were demultiplexed and mapped to the reference genome GRCh38 (build 2020-A, 10X Genomics) using the Space Ranger software v.1.3.0 (10x Genomics) pipeline version 2021.0614.1. Spots were annotated by a specialist breast pathologist using the Loupe v.5.1.0 software (10x Genomics).

### Bioinformatic analysis

For single-nuclei/cell data, read filtering, barcode and UMI counting were performed using Cell Ranger v7.0. High quality barcodes were selected based on the overall UMI distribution using emptyDrops(Lun et al., 2019). All further analyses were run using the Python-based Scanpy(Wolf et al., 2018). To remove low quality cells, we filtered cells with a high fraction of counts from mitochondrial genes (20% or more) indicating stressed or dying cells(Macosko et al., 2015). In addition, genes expressed in less than 20 cells were excluded. Cell by gene count matrices of all samples were concentrated to a single matrix and values log transformed. To account for differences in sequencing depth or cell size UMI counts were normalized using analytical pearson residuals(Lause et al., 2021).

Visium data was analysed using Squidpy (Palla et al., 2022). We used Tangram(Biancalani et al., 2021) to map the annotated snPATHO-Seq annotated data into the spatial transcriptomics data. The top 100 DEG Marker genes in the single nuclei clusters were selected as training genes for Tangram to project snPATHO-Seq to Visium. Then, the normalized cell type probabilities were visualized to obtain sample composition plots.

## Results

### Probe-based chemistry dramatically increases the transcript capture in fixed samples

Sample preservation separates the sampling site and time from the subsequent processing stages. To evaluate the utility of the Fixed RNA Profiling (FRP) kit from 10x Genomics’ as a sample preservation and gene expression profiling general platform, we compared the FRP kit to the conventional Chromium NextGEM Single-cell 3’ kit with v3.1 chemistry. For this evaluation, we used the prostate cancer LnCAP cell line, which was readily available to us. Remarkably, after sequencing depth equalizing, the FRP kit captured 16.7% more UMIs than the v3.1 kit when using the whole transcriptome, however this difference increased to 2.1x more UMIs captured (101 million vs 49 million) when only the genes in the Probe Set are considered (**Figure 1A**). The FRP technology improved the detection of 11624 genes (81.5%) and, furthermore, it was able to detect 83 genes that were undetectable in the v3.1 derived data (Supplementary Table 1). Not surprisingly, most of the genes that were better captured in the fresh sample were ribosomal and mitochondrial genes. The increased UMI capture of informative transcripts in the FRP kit together with the low mitochondrial genes detected was translated in a 99.9% of cells passing the QC filters versus 81.6 % using the v3.1 chemistry (**Supplementary Figure 1**). At matched sequencing depth (24,000 reads/cell), fixed sample showed more UMIs, and genes detected (**Figure 1B**). Detection of low expressed genes remains challenging due to technical dropouts using standard poly-A capture/Template-Switching strategies. To that end, we used a subset of all human transcription factors (TF) to compare the level of detection between FRP and v3.1 kits. Analysis of the top 20 TF including the established prostate cancer marker AR and common oncogene TP53 showed overall better transcript capture from the fixed sample than the fresh sample (**Figure 1C**). A comparative table with all human TF can be explored as Supplementary table 2. Interestingly, cell cycle enrichment analysis showed a define clustered into different cell cycle stages which correlates with the expression of MKI67 for the fixed sample (**Figure 1D**). However, for the fresh sample the clustering was not clear. Taking all together, the FRP kit increased the gene expression information obtained from the cell line by dramatically boosting the capture/detection, reducing the technical dropouts and focusing the sequencing space on informative genes.

**Figure 1.**
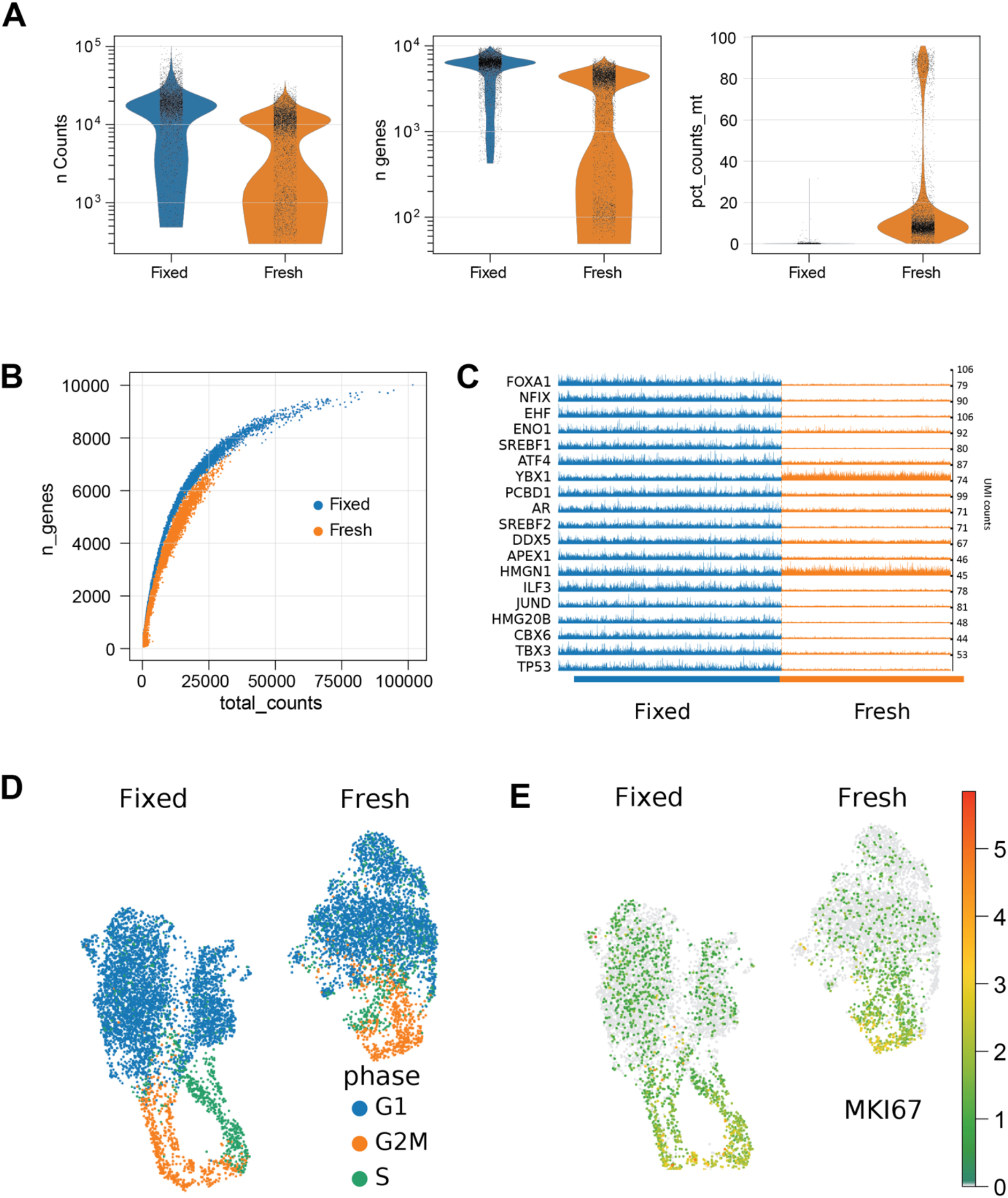
10X Fixed RNA Profiling (FRP) kit validation on a matched pair of fresh-fixed sample **A)** Approximately 5000 cells were analysed using the FRP (Fixed) or the Chromium Single-cell 3’ v3.1 kits respectively. Fastq files were down-sampled to match 24K read/sample in both samples. Basic QC statistics on detected genes, UMI counts, and percentage of mitochondrial genes are shown. **B)** Saturation plot comparing fresh vs. fixed sample. **C)** Tracks plot comparing the detection of the top 20 Transcription factors detected in the sample. **D)** Cell cycle enrichment score calculated for fixed and fresh samples integrated in the same UMAP space. **E)** MKI67 gene expression in fixed and fresh sample. Process data can be accessed as .h5ad including raw counts here <u>Link to Colab</u> (https://github.com/afvallejo/snPATHOSeq/blob/main/FRP_validation_using_LnCaP.ipynb) .

### Unlocking the FFPE archives for single nucleus RNA profiling

FFPE tissue repositories are an enormous resource for the identification of biomarkers, disease pathways, and drug targets. We have optimized a method for efficiently extracting intact nuclei from FFPE tissue sections and perform single nuclei RNA profiling, namely snPATHO-Seq. In brief, the method involves removing the paraffin, performing a partial-to-total enzymatic digestion, and extraction of intact nuclei using a lysis buffer.

We challenged this new method with an archived breast cancer liver metastasis tissue sample collected during autopsy. The autopsy sample was collected at least 4 hours after the death of the patient. Hence, RNA quality in this sample was impacted by both the chemical and physical stresses during FFPE sample preservation and post-mortem RNA degradation. Nonetheless, our method enabled the isolation of high-quality nuclei without the need of a FACS instrument (**Figure 2A**), although FACS using DAPI staining is optional. Purified nuclei are then processed using the FRP kit (10x Genomics) with adjusted cycling conditions (see Methods, **Figure 2B**). A total of 800,000 nuclei in 500 μL volume were available after hybridisation with the gene targeting probes, of which ∼8,000 were used for Chromium instrument, while the remainder were stored and available for future captures if needed further nuclei are needed for analysis. After data processing, we were able to identify 5721 nuclei with a median of 3216 UMI and 1850 genes detected per cell.

**Figure 2.**
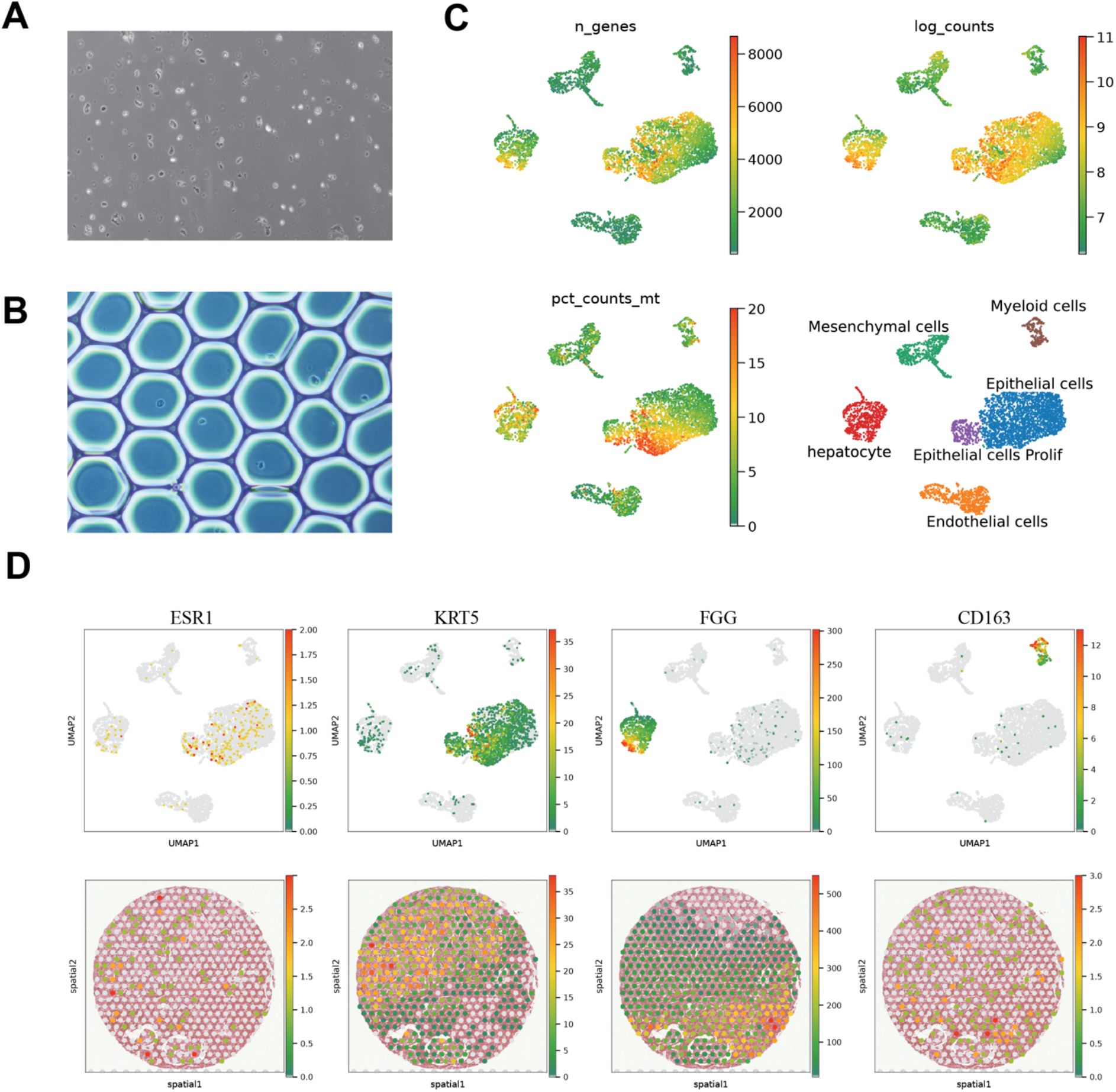
snPATHO-Seq. **A)** representative images of purified nuclei and **B)** Gel Bead-In Emulsion (GEM) from FFPE archive samples showing captured single nuclei. **C)** QC UMAP plots showing average detected genes, log2 number of UMI counts and percentage of mitochondrial genes. Sample was annotated using cell markers in Supplementary figure 2. **D)** Comparison of marker detection in snPATHO-Seq (top) and Visium (bottom) for adjacent samples. Colour scale represent UMI counts/cell.

After normalization, dimension reduction and clustering, we identified 5 major clusters (after grouping epithelial and epithelial proliferative) including a hepatocyte cluster, representing major cell lineages expected to be present in breast cancer liver metastasis (Figure 2C). Interestingly, no clusters for T cell or B cell were identified in the current dataset, though this may be expected as liver metastases are generally considered to be ‘immune deserts’. However, we were able to identify cancer cells, myeloid cells, endothelial and stromal cell clusters. Importantly, we identified a cluster of hepatocytes (**Figure 2C**) which are challenging to profile using conventional, polyA-based, single-cell capturing strategies (Slyper et al., 2020) (**Figure 2C**).

Taking advantage of the flexibility in sample selection of our new method, we validated the findings in single-nuclei data using the Visium spatial transcriptomic technology with matched FFPE samples. While the current Visium data was generated from adjacent samples and only a small region was targeted in the Visium assay, one can imagine applying the snPATHO-Seq and Visium technology to serial sections obtained from the same FFPE block will enable more accurate cell type mapping and cross-evaluation. We evaluated the expression of cell markers in the Visium data and observed consistent expression patterns between data generated by snPATHO-Seq and Visium (**Figure 2D**). Importantly, the expression of T and B cells markers such as CD3D and MS4A1 also appears to be largely missing in the Visium and were not detected in the snPATHO-Seq data (**Figure 2D**). In line with previous literature, the lack of T and B cell clusters likely reflects the biological nature of late-stage metastatic breast cancer with limited immune cell infiltration(Szekely et al., 2018; Zhu et al., 2019).

Furthermore, both the snPATHO-Seq method and the Visium assay are effective tools in resolving intra-tumour heterogeneity. The donor of the metastatic breast cancer tissue sample was firstly diagnosed with ER positive breast cancer which subsequently switched to the triple negative subtype following treatment. While the sample used in the current study was collected as a triple negative breast cancer sample by clinical examination, we identified cancer cells (snPATHO-Seq) and cancer related spots (Visium) with ESR1 expression suggesting the presence of residual luminal breast cancer cells (**Figure 2D**). While further investigation on the molecular nature of these cells and spots is still needed, our current results have demonstrated the sensitivity of the snPATHO-Seq and Visium technologies and their value in resolving tumour heterogeneity.

The FRP and Visium FFPE v1.0 kits use similar probe-based chemistry (not identical probeset); however, the probe sequence and tiling are not the same. To compare the capture rate among both assays we used pseudo-bulk grouping and correlation analysis. Comparison of the common detected genes showed a correlation of 0.9 (**Figure 3A**). In addition, the comparison of all detected genes in both techniques showed an 84.2% overlapping in the gene detection with 5.5% (949 genes) only detected by snPatho-Seq (**Supplementary figure 3**). With the high correlation on gene detection and detection chemistry among both techniques, we explored the potential use of snPATHO-Seq for inform ST sample deconvolution. Specialist breast pathologist annotation was used as ground truth (**Figure 3B**). After cell segmentation, we used Tangram to unbiasedly project snPATHO-Seq to Visium data. Sample deconvolution labels correlated with the marker genes used for annotation (**Figure 2D and 3C**). Moreover, we observed a high correlation in cellular composition between the FRP and Visium datasets. Total epithelial cell (including proliferative) was 68% on the snPATHO-Seq data whereas ST deconvolution detected 70%. We also detected comparable proportions of hepatocytes and stromal cell types between the snPATHO-Seq and Visium data while the proportion of immune infiltration appeared to be low in both datasets (**Figure 3D**).

**Figure 3.**
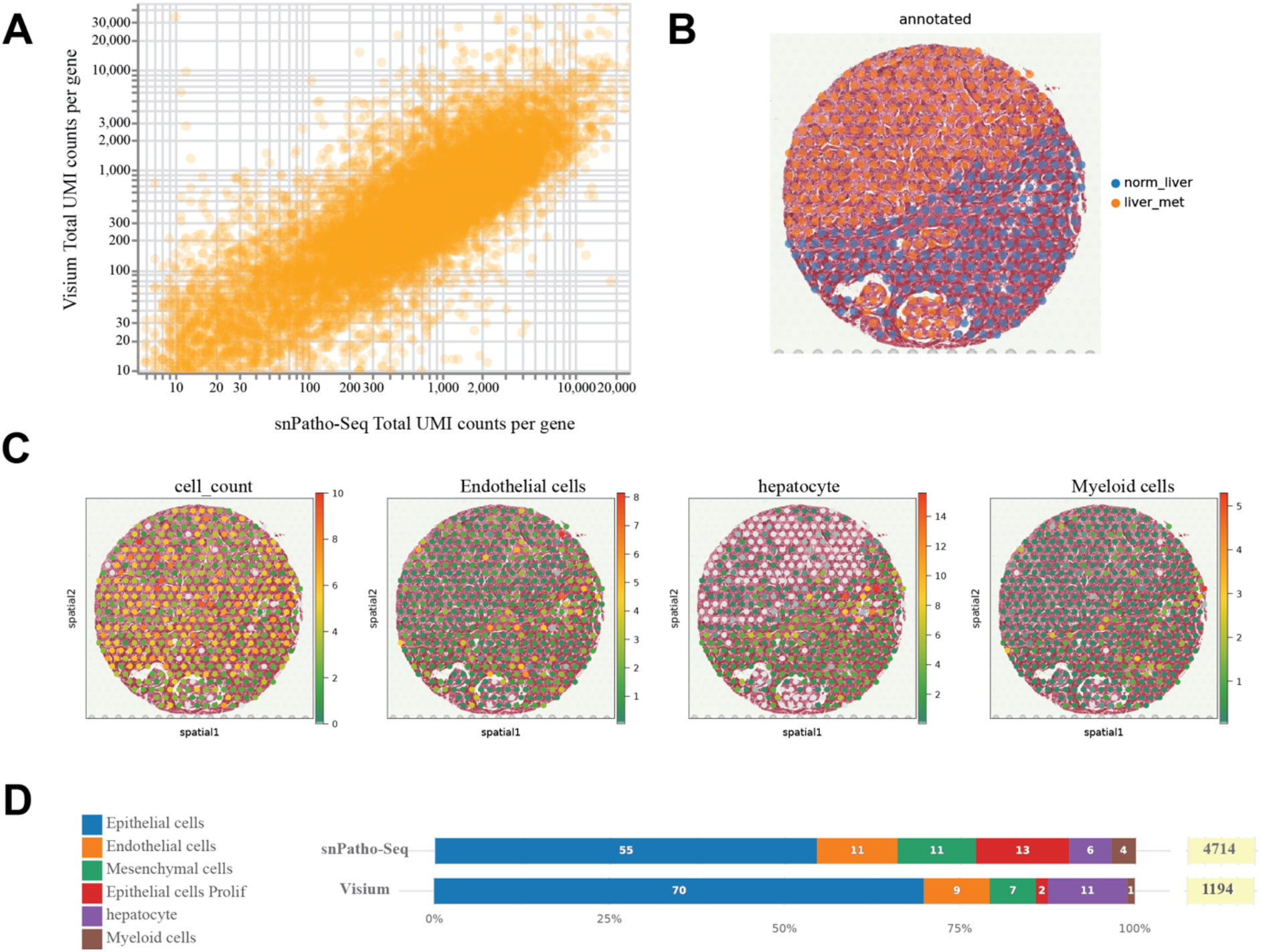
snPATHO-Seq. **A)** Pseudo-Bulk correlation of common genes comparing snPATHO-Seq (X axis) vs Visium (Y axis). Correlation among the samples was 0. **B)** Cancer cell distribution pattern as pe specialist breast pathologist assessment. **C)** snPATHO-Seq projection into Visium data using Tangram. After cell segmentation, the number of cells per dot was estimated. The top 100 DEG Marker genes in the single nuclei clusters were selected as training genes for Tangram to project snPATHO-Seq to Visium **D)** Comparison of sample composition using snPATHO-Seq and the projected composition from the Visium data.

## Supporting information

Supplementary Protocol

## Acknowledgments

This work is supported by The National Health and Medical Research Council (NHMRC) via an Ideas Grant (APP2004774), and the Commonwealth Standard Grant Agreement (4-F26M8TZ). The funders had no role in study design, data collection and analysis, decision to publish or preparation of the manuscript. We thank Jens Durruthy Durruthy, Daniel “Telstra” Dlugolensky, Andrew Kohlway from 10x Genomics and the team at Millennium Sciences for the technical support and for providing access to the Fixed RNA Profiling kit.

## References

Basile, G., Kahraman, S., Dirice, E., Pan, H., Dreyfuss, J. M., & Kulkarni, R. N. (2021). Using single-nucleus RNA-sequencing to interrogate transcriptomic profiles of archived human pancreatic islets. Genome medicine, 13(1), 1–17.

Biancalani, T., Scalia, G., Buffoni, L., Avasthi, R., Lu, Z., Sanger, A., Tokcan, N., Vanderburg, C. R., Segerstolpe, Å., & Zhang, M. (2021). Deep learning and alignment of spatially resolved single-cell transcriptomes with Tangram. Nature methods, 18(11), 1352–1362.

Fischer, F., Roenneberg, S., Graner, L., Schlenker, F., Zengerle, R., Theis, F. J., Schmidt-Weber, C. B., Biedermann, T., Lauffer, F., & Garzorz-Stark, N. (2022). Gene expression based molecular test proves clinical validity as diagnostic aid for the differential diagnosis of psoriasis and eczema in formalin fixed and paraffin embedded tissue. medRxiv.

Gilbert, M. T. P., Haselkorn, T., Bunce, M., Sanchez, J. J., Lucas, S. B., Jewell, L. D., Marck, E. V., & Worobey, M. (2007). The isolation of nucleic acids from fixed, paraffin-embedded tissues–which methods are useful when? PloS one, 2(6), e537.

Hester, S. D., Bhat, V., Chorley, B. N., Carswell, G., Jones, W., Wehmas, L. C., & Wood, C. E. (2016). Editor’s highlight: dose–response analysis of RNA-Seq profiles in archival formalin-fixed paraffin-embedded samples. Toxicological Sciences, 154(2), 202–213.

Jang, J. S., Holicky, E., Lau, J., McDonough, S., Mutawe, M., Koster, M. J., Warrington, K. J., & Cuninngham, J. M. (2021). Application of the 3’ mRNA-Seq using unique molecular identifiers in highly degraded RNA derived from formalin-fixed, paraffin-embedded tissue. BMC genomics, 22(1), 1–9.

Klopfleisch, R., Weiss, A., & Gruber, A. (2011). Excavation of a buried treasure–DNA, mRNA, miRNA and protein analysis in formalin fixed, paraffin embedded tissues. Histology and histopathology, Vol. 26, nº6 (2011).

Lause, J., Berens, P., & Kobak, D. (2021). Analytic Pearson residuals for normalization of singlecell RNA-seq UMI data. Genome biology, 22(1), 1–20.

Lun, A. T. L., Riesenfeld, S., Andrews, T., Dao, T. P., Gomes, T., participants in the 1st Human Cell Atlas, J., & Marioni, J. C. (2019). EmptyDrops: distinguishing cells from empty droplets in droplet-based single-cell RNA sequencing data. Genome Biol, 20(1), 63. https://doi.org/10.1186/s13059-019-1662-y

Macosko, E. Z., Basu, A., Satija, R., Nemesh, J., Shekhar, K., Goldman, M., Tirosh, I., Bialas, A. R., Kamitaki, N., Martersteck, E. M., Trombetta, J. J., Weitz, D. A., Sanes, J. R., Shalek, A. K., Regev, A., & McCarroll, S. A. (2015). Highly Parallel Genome-wide Expression Profiling of Individual Cells Using Nanoliter Droplets. Cell, 161(5), 1202–1214. https://doi.org/10.1016/j.cell.2015.05.002

Martelotto, L. G., Baslan, T., Kendall, J., Geyer, F. C., Burke, K. A., Spraggon, L., Piscuoglio, S., Chadalavada, K., Nanjangud, G., & Ng, C. K. (2017). Whole-genome single-cell copy number profiling from formalin-fixed paraffin-embedded samples. Nature medicine, 23(3), 376–385.

Mehine, M., Khamaiseh, S., Ahvenainen, T., Heikkinen, T., Äyräväinen, A., Pakarinen, P., Härkki, P., Pasanen, A., Bützow, R., & Vahteristo, P. (2020). 3’ RNA sequencing accurately classifies formalin-fixed paraffin-embedded uterine leiomyomas. Cancers, 12(12), 3839.

Palla, G., Spitzer, H., Klein, M., Fischer, D., Schaar, A. C., Kuemmerle, L. B., Rybakov, S., Ibarra, I. L., Holmberg, O., & Virshup, I. (2022). Squidpy: a scalable framework for spatial omics analysis. Nature methods, 19(2), 171–178.

Slyper, M., Porter, C., Ashenberg, O., Waldman, J., Drokhlyansky, E., Wakiro, I., Smillie, C., Smith-Rosario, G., Wu, J., & Dionne, D. (2020). A single-cell and single-nucleus RNA-Seq toolbox for fresh and frozen human tumors. Nature medicine, 26(5), 792–802.

Szekely, B., Bossuyt, V., Li, X., Wali, V., Patwardhan, G., Frederick, C., Silber, A., Park, T., Harigopal, M., & Pelekanou, V. (2018). Immunological differences between primary and metastatic breast cancer. Annals of Oncology, 29(11), 2232–2239.

Wolf, F. A., Angerer, P., & Theis, F. J. (2018). SCANPY: large-scale single-cell gene expression data analysis. Genome Biol, 19(1), 15. https://doi.org/10.1186/s13059-017-1382-0

Zhu, L., Narloch, J. L., Onkar, S., Joy, M., Broadwater, G., Luedke, C., Hall, A., Kim, R., Pogue-Geile, K., & Sammons, S. (2019). Metastatic breast cancers have reduced immune cell recruitment but harbor increased macrophages relative to their matched primary tumors. Journal for immunotherapy of cancer, 7(1), 1–10.

