## Supplementary figures and images for "snPATHO-seq: unlocking the FFPE archives for single nucleus RNA profiling"

### Supplementary Figure 1

#### Fresh

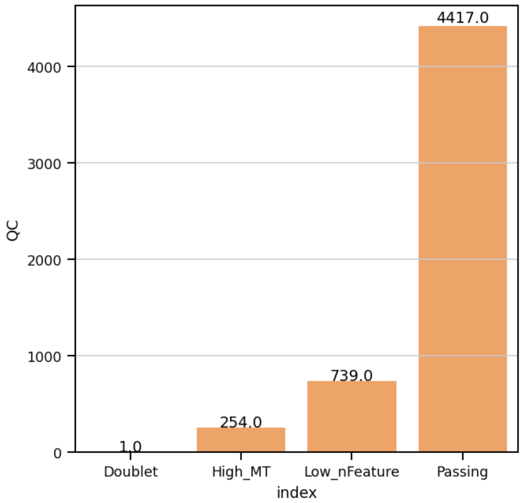


**18% of cells are removed**

#### Fixed

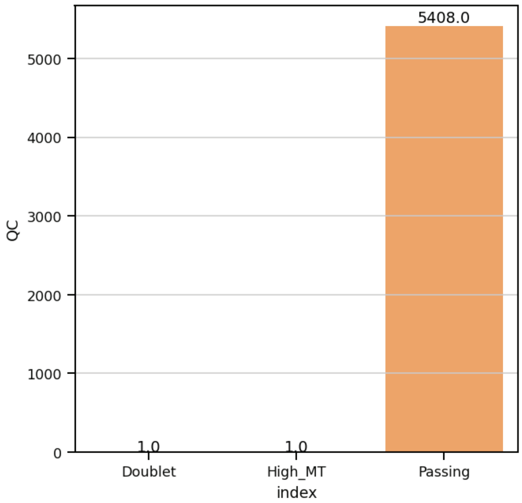

### Supplementary Figure 2

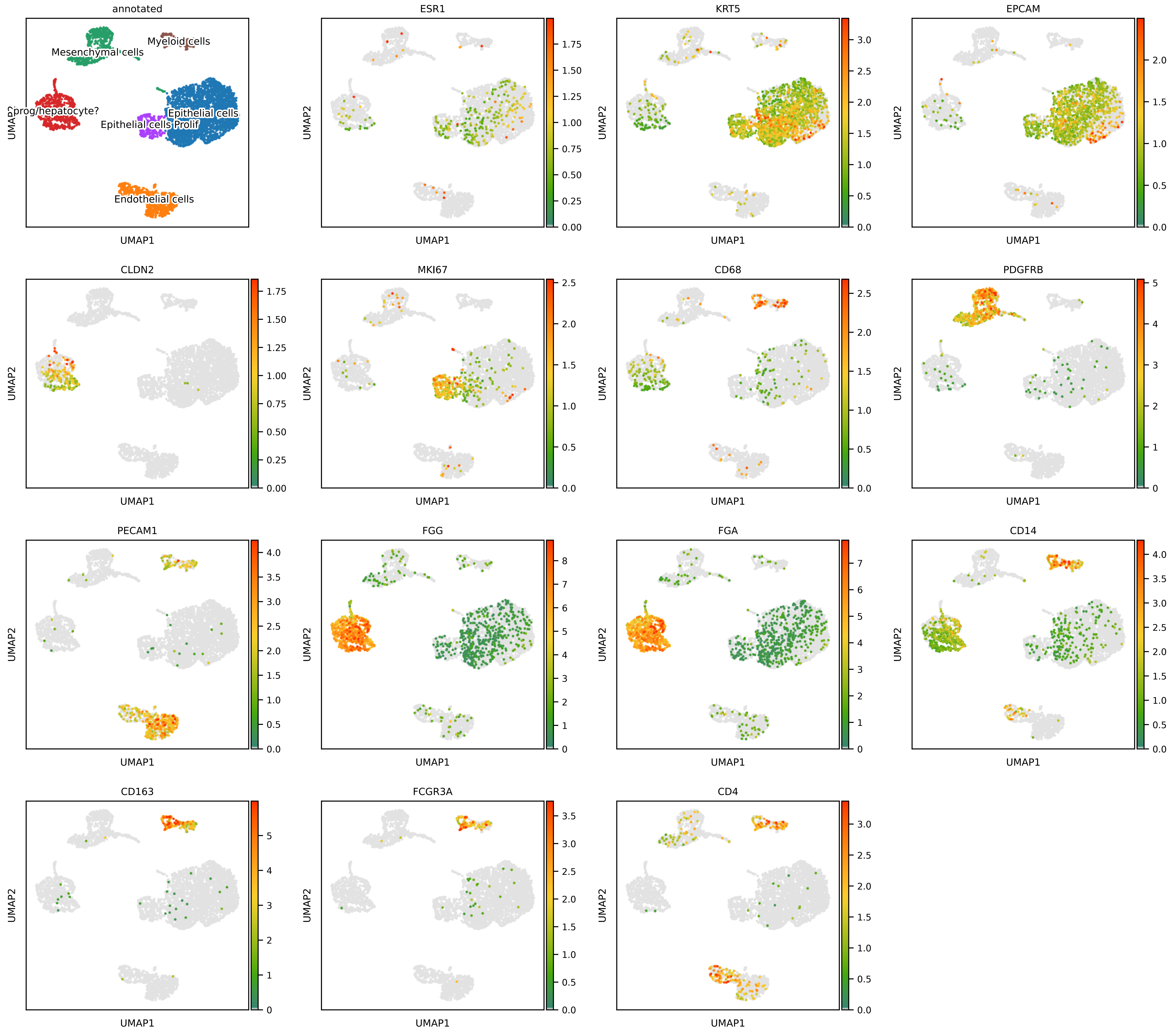

### Supplementary Figure 3

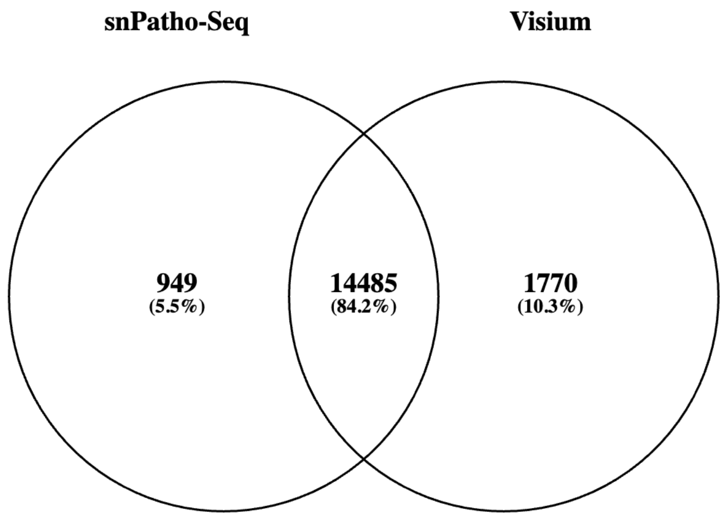
