## Supplementary Protocol for "snPATHO-seq: unlocking the FFPE archives for single nucleus RNA profiling"

### Nuclei suspension preparation for snPATHO-Seq

***Reagents and consumables***

Ethanol
Xylene
Nuclease Free water
1x Phosphate Buffer Saline (PBS, $Ca^{2+}$ and $Mg^{2+}$ free)
Liberase TM (Roche)
Collagenase D (Roche)
$MgCl_2$
10% BSA
EZ Lysis Buffer (Sigma)
10% BSA (MACS® BSA Stock Solution, Miltenyi)
Glycerol 50%
RNAse Inhibitor (RiboLock from Thermo or RNA Protector from Roche)
(Optional) 4',6-Diamidino-2-Phenylindole, Dihydrochloride (DAPI, ThermoFisher)
40 µm and 70 µm pluriStrainer filters (pluriSelect) (MACS SmartStrainers are also possible) is also possible)
25 G needle

***Equipment***

Thermomixer with adjustable shaking (Eppendorf)
Swinging bucket refrigerated centrifuge

**Procedure**

***Nuclei suspension preparation***

1. Cut up to 2 >25 µm-thick sections (punches are also possible) and place it in 1.5 mL Eppendorf tube. Store dry at 4°C if not used immediately. To keep it dry, you may use the cylinder containing silica beads that comes with 10x Genomics chips.
2. Wash sections or punches three times with 1 mL Xylene <u>for 10'</u> to remove the paraffin, rehydrate in sequential 1' of 1 mL ethanol immersions (2× 100%, followed by 1× 70%, 50% and 30% ethanol). IMPORTANT: make sure paraffine is fully removed or digestion will be suboptimal.
3. Wash 3 times (2× 1 mL wash and 1× 800 µL final wash) with 1× PBS + 0.5 mM $CaCl_2$
4. Remove as much volume as possible, and digest tissue for 5-60'(*) at 37°C(**) in 1 mL of 1× PBS + 0.5 mM $CaCl_2$ + 250 µg/mL Liberase +2.5 mg/mL of Collagenase +1 U/µL RNAse Inhibitor. (*) NOTE: some blocks require longer digestion time, so inspect visually and help dissociation by pipetting up and down with a P1000 pipette. (**) Incubation is done in a Thermomixer 800 rpm. IMPORTANT: dissociation does not need to be complete; the objective here is to loosen up the material to facilitate the nuclei release. Dissociation completeness varies from block to block.

snPATHO-Seq

5. Next, add 400 µL of Ez Lysis Buffer to the sample, mix by inverting 5× and centrifuge for 5' at 850xRCF at 4°C.
6. Resuspend the pellets (released nucs and undigested tissue) in 250 µL Ez Lysis buffer + 2% BSA + 1 U/µL RNAse Inhibitor and homogenize the sample using a douncer/pestle by stroking 10-20 times (or as needed).
7. After homogenization add 750 uL Ez Lysis buffer + 2% BSA + 1 U/µL RNAse Inhibitor and continue disaggregating by pipetting using a P1000 pipette (10 times). Incubate on ice for 10'. At 5' mark pipette up and down using a P1000 pipette (10 times). Optional but very useful when possible: if the dissociation and disaggregation look almost complete (i.e., only very small chunks of undigested tissue or fat are visible to the naked eye) gently pass the sample through a 25 G needle for 20 times (don't make foam!). It is key to ensure that large chunks are gone before passing though needle. If large chunks or fat remain the needle will definitely block; so skip this step. This optional step will increase the nuclei release.
8. Pass sample through a 70 µm PluriStrainer filter (not Flowmi!) and centrifuge the flowthrough for 5' at 850xRCF at 4°C and wash nuclei suspensions once more with 800 µL of EzLysis buffer + 2% BSA + 1 U/µL RNAse Inhibitor.
9. Pellet the nuclei for 5' at 850xRCF at 4°C and resuspend in 500 µL of 1x Fix & Perm Buffer (PN-2000517) for 1 h at RT. Then, pass through a 40 µm PluriStrainer filter (not Flowmi!) and then pellet nuclei for 5' at 850xRCF at 4°C.
10. Wash nuclei twice PBS 0.5x + 0.02% BSA and resuspend in 500-1000 µL of PBS 0.5x + 0.02% BSA (variable depending on pellet size). Count using Luna-FX7 or similar based on dual-fluorescence such as AO/PI.
11. Rest on wet ice for immediate FACS cytometry analysis/sorting or proceed to the Chromium X run using Chromium Fix RNA Profiling (10x Genomics) following the user guide for singleplexed or multiplexed samples accordingly. Alternatively, supplement the sample with 0.1x volume of Enhancer solution (10x Genomics) + 10% Glycerol, rest on ice for 10' and cryopreserve at −80°C. Notes: cycling conditions for Index PCR might need to be optimized per sample to obtain a final library that falls within ~50-200 nM. For nuclei derived from FFPE blocks, we typically use 1-2 additional cycles during indexing to start with. If the library does not reach the recommended range but the Bionalyzer/Fragment Analyzer/Tapestation traces look as expected (single peak at ~265 bp), then do not add additional cycles. If you see signs of under/over amplification in the traces, then adjust cycling accordingly.
12. After storage, centrifuge sample for 5' at 850xRCF at room temperature. Remove the supernatant without disturbing the pellet and continue from step 2.1a or even 2.1m (as recommended by 10x Genomics) of 2.1 Post-Hybridization Wash.

snPATHO-Seq

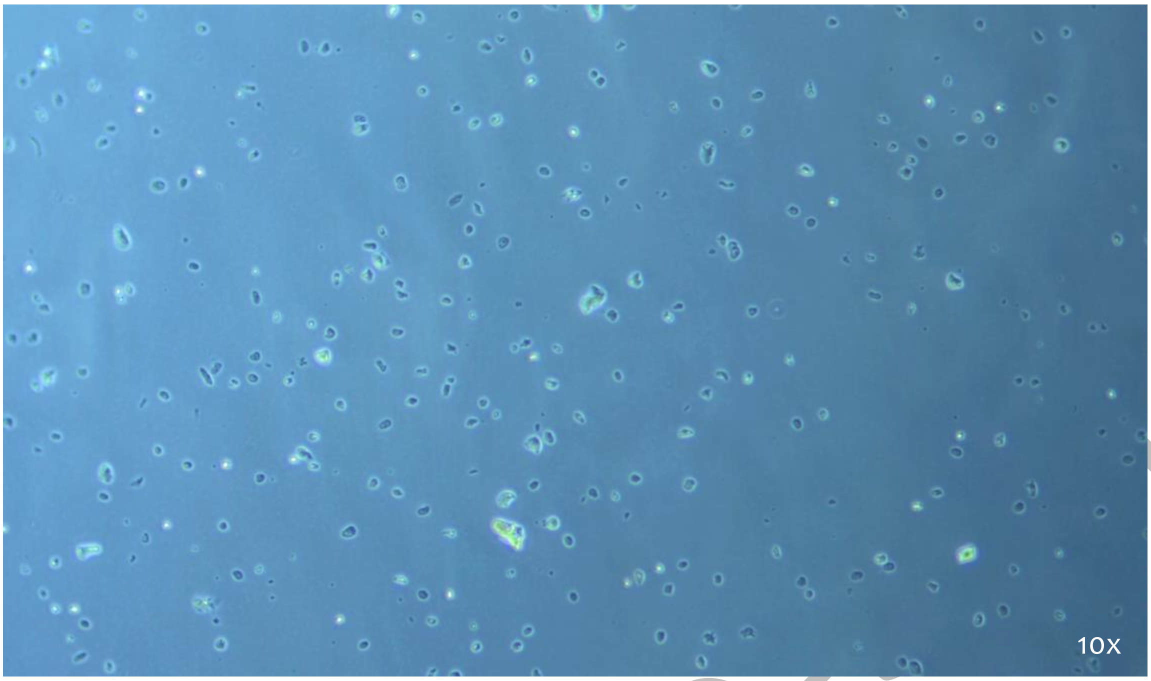
10x

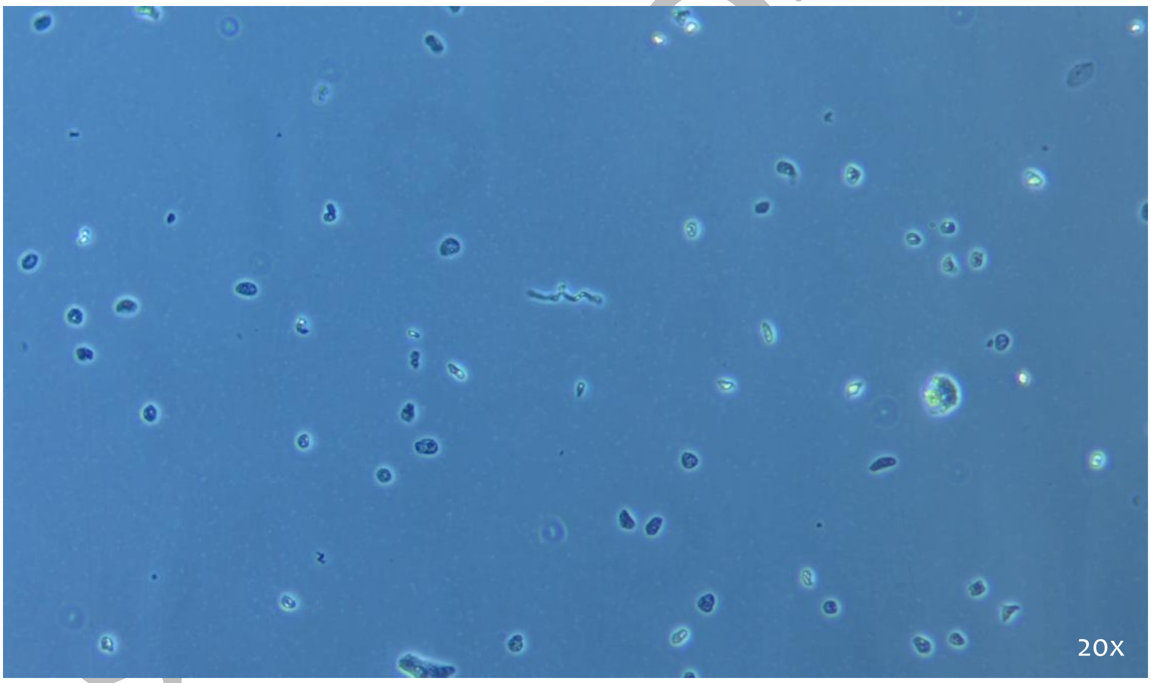
20x
